# X-ray Crystallographic and Hydrogen Deuterium Exchange Studies Confirm Alternate Kinetic Models for Homolog Insulin Monomers

**DOI:** 10.1101/2024.04.16.589767

**Authors:** Esra Ayan, Miray Türk, Özge Tatlı, Sevginur Bostan, Elek Telek, Baran Dingiloğlu, Büşra Züleyha Dogan, Muhammed Ikbal Alp, Ahmet Katı, Gizem Dinler-Doğanay, Hasan DeMirci

## Abstract

Despite the crucial role of various insulin analogs in achieving satisfactory glycemic control, a comprehensive understanding of their in-solution dynamic mechanisms still holds the potential to further optimize rapid insulin analogs, thus significantly improving the well-being of individuals with Type 1 Diabetes. Here, we employed hydrogen-deuterium exchange mass spectrometry to decipher the molecular dynamics of newly modified and functional insulin analog. A comparative analysis of H/D dynamics demonstrated that the modified insulin exchanges deuterium atoms faster and more extensively than the intact insulin aspart. Additionally, we present new insights derived from our 2.5 Å resolution X-ray crystal structure of modified hexamer insulin analog at ambient temperature. Furthermore, we obtained a distinctive side-chain conformation of the Asn3 residue on the B chain (AsnB3) by operating a comparative analysis with a previously available cryogenic rapid-acting insulin structure (PDB_ID: 4GBN). The experimental conclusions have demonstrated compatibility with modified insulin’s distinct cellular activity, comparably to aspart. Additionally, the hybrid structural approach combined with computational analysis employed in this study provides novel insight into the structural dynamics of newly modified and functional insulin vs insulin aspart monomeric entities. It allows further molecular understanding of intermolecular interrelations driving dissociation kinetics and, therefore, a fast action mechanism.

## Introduction

Insulin plays a critical role in regulating cellular glucose uptake. Deficiency or insensitivity to insulin leads to diabetes mellitus, a prevalent and severe metabolic disorder. Treatment typically involves administering insulin and derivatives, which are integral interventions (Ayan, 2022). Insulin is primarily hexameric and dimeric during production, delivery, and circulation. Nonetheless, when attaching to its receptor, it favors a monomeric conformation (DeFelippis et al., 2001), distinct from its structure in solution as marked in NMR and X-ray crystallographic studies (Weis et al., 2018; Scapin et al., 2018). Environmental conditions and alterations in sequence that promote the monomeric state boost fibrillation (Brange et al., 1997) and degradation (Brange et al., 1993), ultimately compromising the stability of therapeutic formulations. Therefore, there is considerable interest within medicine, biophysics, and structural biology in comprehending the structural configurations accessible to the insulin monomer under physiological circumstances. Structural analyses of the fully active insulin monomer (Bocian et al., 2008) largely favor the hexameric form’s T-state identified in crystallographic studies (Baker et al., 1988). Research studying amide hydrogen exchange for an analog present as a monomer in solution reveals limited protection at the A-chain N-terminus, the B-chain N-terminus, and the B-chain C-terminus (Hua et al., 2011). The exchange followed at the relevant locations proposes that those parts examine solvent-exposed conditions, though the specific nature of these states still needs to be fully comprehended. Experimental examination of the structural ensemble of the wild-type monomer is challenging due to both fibrillation and oligomerization occurring from micromolar to millimolar (Weiss et al., 1991). Assorted experimental structural monomers primarily stem from investigations involving sequences with substitutions or studies conducted under in solution that deviate greatly from physiological parameters (e.g., pH > 8), potentially impacting the extent of the disorder (Tokumoto et al., 2006; Fredericq, 1953). More comprehensive all-atom molecular dynamics inquiries need to concentrate on the insulin monomer. Nevertheless, simulations present a direct avenue for investigating microscopic structures and the fundamental stabilizing forces. Nanosecond-scale unbiased simulations were conducted on the porcine insulin monomer, initiating from the T state depicted in the hexameric crystal structure (Baker et al., 1988). These simulations unveiled disorder within the N- and C-termini of the B-chain, while no disorder was observed in the AN-helix. The root-mean-square deviation from the T state proposes considerable unfolding, though with restricted structural elucidation. Additionally, bias-exchange metadynamics simulations on porcine insulin at low pH and elevated temperature yielded a varied ensemble predominantly consisting of fully unfolded states (Singh et al., 2017). The structural characterization of a protein containing disordered regions poses experimental difficulties due to the rapid interconversion of conformations within picosecond to millisecond lifetimes (Bonomi et al., 2017). While hydrogen/deuterium exchange and mass spectrometry (HDX/MS) represents a valuable approach for gaining insights into protein conformation and disordered structural dynamics (Spolaore et al., 2023; Oliva, 2024), previous studies have focused on conducting H/D reactions on oligomeric insulins rather than monomeric conformer (Nakazawa et al., 2012;2012). This study attempted to elucidate the underlying mechanism governing the dynamic changes in the monomer of less-stable insulin analog compared to its counterpart, insulin aspart (IAsp). This was achieved by introducing a mutation in the 22^nd^ residue on the B chain, replacing arginine with a more flexible lysine residue together with a substitution at the 28^th^ residue on chain B from proline to aspartic acid, and investigating its impact on protein stability and electrostatic interactions (hereafter referred to as INSv). The model system employed for this study was IAsp (Simpson and Spencer, 1999). Once substitution, the activity of the INSv monomer in the cells has been confirmed over calcium imaging, serving the IAsp monomer as a control. Subsequently, the stability of the INSv monomer was assessed using the differential scanning calorimetry (DSC) technique, comparing with IAsp monomer stability. Hydrogen-deuterium exchange mass spectrometry (HDX-MS) analysis explored these monomers’ intrinsic and distinct dynamics. The findings from this experimental approach were further validated computationally by employing Gaussian Network Model (GNM) analysis. Finally, a 2.5 Å resolution X-ray crystal structure of its hexamer was determined through the newly developed multi-well X-ray crystallography technique of Turkish Light Source conducted at ambient temperature, elucidating the conformational differences between the hexamers of both INSv and IAsp.

## Methods

### Production of recombinant LysB22-AspB28 double mutant (INSv)

The recombinant INSv gene was cloned to the pET28a(+) vector and was transformed into the *E. coli* BL21 Rosetta-2 strain for overexpression. Bacterial cell cultures, each containing the recombinant INSv protein genes, were cultivated independently in 6 liters of either standard LB-Miller media or Terrific Broth (TB), supplemented with the final concentration of 35 μg/mL chloramphenicol and 50 μg/mL kanamycin at 37°C. Cultures were agitated by using a New Brunswick Innova 4430R shaker at 110 rpm until they reached an optical density at 600 nm (OD_600_) of approximately 0.7 for each culture. Recombinant protein expression was induced with Isopropyl b-D-1-thiogalactopyranoside (IPTG) at a final concentration of 0.4 mM. The incubation for protein overexpression was conducted at 37°C for 4 hours. Subsequently, cells were harvested at 4°C using a Beckman Allegra 15R desktop centrifuge at 3500 rpm for 20 minutes. Verification of protein expression was conducted through precast TGX-mini protean gradient SDS-PAGE from BioRad.

### The solubilization process of INSv

For the partial purification and complete solubilization of inclusion bodies, cell pellets (1 gram) containing INSv proteins were individually resuspended in 10 mL of lysis buffer containing 25 mM Tris-HCl, pH 8.0, 5 mM Ethylenediaminetetraacetic acid (EDTA) and subsequently homogenized. Bacterial cells were lysed via sonication, and the resulting cell debris underwent low-speed centrifugation (6000 × g, 4°C, 7 minutes). The obtained pellets were subjected to suspension in wash buffer A containing 25 mM Tris-HCl, pH 8.0, 5 mM EDTA supplemented with 0.1% (v/v) Triton X-100 and 1 M urea, followed by wash buffer B containing 25 mM Tris and 2 M urea. The suspension was sonicated for 30 seconds in an ice bath and then centrifuged (8000 × g, 4°C, 20 minutes). To further isolate inclusion bodies, the debris was resuspended at 0.1 g/mL in binding buffer containing 25 mM Tris-HCl, pH 8.4, and 5 mM 2-mercaptoethanol (BME) for INSv, which contained 8 M urea. Centrifugation at 10,000 × g for 30 minutes at 4°C allowed recovery of inclusion bodies containing the fusion proteins, and the resulting supernatants were filtered through a 0.2-micron Millipore filter.

### Purification, refolding, and tryptic digestion process of INSv

Sulfitolysis of INSv was conducted at 25°C for 4 hours using 200 mM sodium sulfite (NaLSOL) and 20 mM sodium tetrathionate (Na_2_S_4_O_6_). Subsequently, the manual gravity Ni-NTA affinity chromatography was employed, involving column equilibration with one column volume of a buffer containing 25 mM Tris-HCl, pH 8.0, 5 mM BME, and 8 M urea and elution with a buffer containing 25 mM Tris-HCl, pH 8.0, 5 mM BME, 8 M urea, and 250 mM imidazole. The protein sample purity was assessed at each stage through 20% SDS-PAGE gel analysis and Coomassie staining. High-purity fractions were collected and filtered through a 0.2-micron Millipore filter. The INSv refolding process is completed by using standard published protocols (Khosravi et al., 2022; Kim et al., 2015; Min et al., 2011). Purified proinsulin (0.5 mg/mL) underwent dialysis against 25 mM Tris-HCl pH 9.00 with 8 M urea and 0.5 mM EDTA at 4°C. After centrifugation (10.000 × g, 4°C, 30 minutes) to remove potential aggregates, the resulting mixture was dialyzed in a refolding solution with 0.1 M Gly/NaOH at pH 10.5, 0.5 M urea, 0.5 mM EDTA, 5% glycerol, and stirred at 15°C overnight. Refolded INSv was further dialyzed against 25 mM Tris-HCl, pH 8.0, with 5% glycerol for 18 hours. Proinsulin underwent tryptic cleavage using trypsin 1:1000 (v/v) for 4 hours at 30°C. Following the proteolytic cleavage, pH precipitation was employed by adding 20 mM citric acid to the protein sample. Upon precipitation, INSv was centrifuged at 3500 rpm at 4°C for 5 minutes, and the resulting pellet was dissolved in 20 mM citric acid to quench trypsin activity and subsequent removal of trypsin. After solubilization, INSv was filtered through a 0.2 μM pore diameter filter and subjected to size exclusion chromatography (SEC) using a 20 mM citric acid buffer. High-purity fractions were collected, and pH precipitation was performed using 1 M Tris, pH 10.0. The INSv pellet was then solubilized using 25 mM Tris-HCl, pH 9.0 buffer, and the concentration was adjusted to 1000 µg/mL for HDX-MS analysis.

### Monomerization of insulin aspart (IAsp)

Precipitation through pH adjustment and subsequent solubilization in 25 mM Tris-HCl pH 9.0 buffer were employed to convert IAsp (NovoRapid) from a hexameric to a monomeric state (Tokumoto et al., 2006; Fredericq, 1953). Using a 10K amicon ultrafiltration unit, a buffer exchange was performed ten times with 25 mM Tris-HCl pH 9.0. The resulting monomeric form of NovoRapid was then adjusted to a 1000 µg/mL concentration for HDX-MS analysis.

### INSv and IAsp-induced in-vitro activity experiments

#### Primary hippocampal cell culture

Newborn mice aged 0-3 days were decapitated following ethical guidelines. Their brains were extracted in a sterile laminar flow cabinet and immediately immersed in a Hibernate-A (Gibco-A1247501) solution, including 1% Antibiotic and 1% Glutamax. The hippocampi were carefully dissected under a stereomicroscope and placed in an enzymatic solution containing L-15 (Sigma L5520), 1% antibiotics, 1% Glutamax, 2% B27, 1% papain, and 1% DNAse, and maintained at 4°C for 45 minutes. Once performing the enzymatic treatment, the hippocampal tissues were mechanically triturated using a series of three glass Pasteur pipettes with sequentially decreasing tip diameters. The resulting cellular suspension was then transferred to an enzyme inhibition medium containing L-15 solution, 1% antibiotics, 1% Glutamax, 2% B27, and 10% FBS and incubated at 4°C for 15 minutes. The cells suspended in the enzyme inhibition medium were centrifuged at 1000 rpm for 5 minutes. The cells were resuspended with Neurobasal-A containing 1% antibiotics, 1% Glutamax, and 2% B27 and seeded into cell culture dishes at a density of 15,000–20,000 cells/mm². After the initial seeding, the cells were incubated at 37°C in an atmosphere containing 5% CO2 for 2 hours. To maintain optimal conditions, the total volume in the culture dish was adjusted to 1500 μL with additional maintenance medium. GCaMP plasmid (Addgene plasmid #100843) (Chen et al., 2013) was transduced to the cells using an adeno-associated virus (AAV) known for its efficacy in neuronal systems. Half of the medium was replenished every three days. Experimental procedures were initiated on day 21, after a significant increase in GCaMP expression and following the maturation of the neuronal network.

#### Calcium imaging of INSv and IAsp-induced activities

Sequential action potentials trigger a substantial influx of calcium ions into the cell. Upon entry, these calcium ions interact with the binding domains of the GCaMP protein, inducing conformational alterations within the protein structure. These structural changes activate the green fluorescent protein (EGFP) linked to GCaMP, enhancing fluorescence intensity. To visualize this fluorescence, the EGFP must be excited using blue light at a wavelength of 488 nm, leading to the emission of fluorescent green light at a wavelength of approximately 515 nm. The calcium activities within hippocampal neuronal networks were monitored and visualized using a fluorescent microscope equipped with a connected camera. Imaging was conducted using 10x objectives. Images were captured at a frequency of 1 frame per second and subsequently compiled into video recordings. The video recordings were analyzed using Zen Blue (Zeiss) software, wherein selected cell bodies were designated as regions of interest (ROIs). The average light intensity variation within these areas was calculated for every frame. The results were further processed and analyzed using Excel (Microsoft) and Prism 9 (GraphPad Software, La Jolla, CA, USA). Temporal variations in light intensity observed within the selected cell bodies were graphically represented over time. The quantification of active cells was conducted through manual enumeration.

### Higher-order insulin monomers defined through HDX-MS analysis

HDX-MS analysis has been performed at five different time points ranging from 12 s to 24 h. Accordingly, the heat maps were generated, and the structural features based on deuterium uptake rate and electrostatic and hydrophobic potentials were visualized by *PyMOL* (DeLano, 2022). Purified INSv and IAsp were diluted in the equilibration buffer containing 50 mM Tris-HCl; pH 9.5. For labeling, 2.5 μl (≥ 35 pmol) of each sample was incubated with 37.5 μl labeling buffer containing 50 mM Tris-HCl in D_2_O; pD 9.5; pH 9.1 at room temperature for 0, 0.2, 2, 20, 200, or 1440 mins incubation periods. Labeled samples were quenched with 40 μl pre-chilled quench buffer containing 50 mM Tris-HCl, 3M guanidine hydrochloride (GnHCl), 100 mM tris(2-carboxyethyl)phosphine (TCEP); pH 2.3, then the sample was incubated on ice to inhibit deuterium back-exchange. All labeling time points were subject to triplicate analyses. t_0_ samples were used as reference. Relative Fractional Uptake graphs were generated by GraphPad Prism 8.1 according to uptake data obtained from *DynamX* Software (Waters Corp., 176016027). Deuterated samples were digested using The Enzymate BEH Pepsin Column (Waters Corp., Manchester, UK) at 20°C. Upon 3 minutes of transferring digested peptides at Waters BEH C18 VanGuard trapping pre-column (flow rate 70 μl/min in 0.2% formic acid), proteins underwent online pepsin digestion and liquid chromatography/mass spectrometry (LC/MS) using the nanoACQUITY UPLC system and the Synapt G2-Si Q-Tof mass spectrometer from Waters Corp. (Milford, MA, USA). Peptide separation conditions included a 6-minute gradient with 0.2% formic acid, 35% acetonitrile, maintained at 0 °C, and a 40 μl/min flow rate. MS data were gathered using an MS^E^ method in resolution mode with an extended range enabled to avoid detector saturation and maintain peak morphologies (Bonn et al., 2010). MS was initially calibrated with [Glu1]-Fibrinopeptide B human (Glu-fib) peptide, and MS data were gathered with lock mass correction using Glu-fib peptide in positive polarity. Peptide identification was performed by *ProteinLynx Global Server* (PLGS, 3.0.3, Waters), and identified peptides were filtered by *DynamX* Software (Waters Corp., 176016027.) according to the following parameters: minimum intensity of 800, the minimum sequence length of 3, maximum sequence length of 20, minimum products of 2, minimum products of per amino acid of 0.1, minimum score of 6, maximum MH+ error threshold of 0 ppm. Only high-quality peptides were used for analyses. Deuterium uptake was calculated by an increase in mass/charge ratio compared to t_0_. *DynamX* Software generated the coverage map and heat map.

### Gaussian Network Model with all C*α* in the INSv and IAsp monomers

We conducted normal mode analysis using ProDy (Bakan et al., 2011). Specifically, we employed the Gaussian Network Model (GNM) analysis for the newly determined INSv structure at ambient temperature and a previously resolved structure at cryogenic conditions (PDB ID: 4GBN). Contact maps were defined based on all Cα atoms, utilizing a cutoff distance of 8.0 Å. N − 1 nonzero normal modes were derived, where N represents the total number of Cα atoms. For each structure, we calculated squared fluctuations over the weighted 10 slowest and 10 fastest modes, enabling a comparison of their global and local motions, respectively.

### Differential Scanning Calorimetry of monomer INSv and IAsp analogs

Differential scanning calorimetry (DSC) was conducted to assess the thermal stability of IAsp and INSv insulin samples at a 2 mg/ml concentration, employing a SETARAM Micro DSC-III calorimeter (Jandel Scientific). The measurements were carried out within the 20 to 100°C temperature range, employing a heating rate of 0.3 K/min. Precision balancing, with an accuracy of ± 0.05 mg, was employed for the IAsp and INSv samples and the reference (20 mM citric acid buffer solution) to mitigate corrections involving the heat capacity of the vessels. A secondary thermal scan of the denatured sample was performed for baseline correction. Analysis of the melting temperature (T_m_), calculation of calorimetric enthalpy (ΔH_cal_), and determination of the change in entropy (ΔS) were executed using the OriginPro® 2022b software (OriginLab Corporation, Northampton, USA).

### Hexamerization and crystallization of INSv insulin analog

The transformation of the INSv sample from its monomeric state to the zinc-bound form involved a zinc precipitation step using a solution comprising 2.4 M NaCl, 100 mM citric acid, and 6 mM ZnCl_2_. Subsequently, solubilization was achieved by using 25 mM Tris-HCl solution at pH 8.0. The zinc-bound INSv samples were then screened for crystallization using the oil technique using the sitting drop vapor diffusion micro-batch. Approximately 3000 commercially available sparse matrix and grid screen crystallization conditions were employed for the initial crystallization screening (Durdagi et al., 2020) In a 72-well Terasaki plate (Cat#654,180, Greiner Bio-One, Austria), equal volumes of crystallization conditions (v/v, 0.83 μl) were mixed in a 1:1 ratio with a 5 mg/ml INSv solution. Each well was covered with 16.6 µl of paraffin oil (Cat#ZS.100510.5000, ZAG Kimya, Türkiye) and then incubated at 4°C. Crystal formation in the wells of the Terasaki plates was observed using a compound light microscope.

#### Sample delivery and data collection

Data collection was performed using Rigaku’s XtaLAB Synergy Flow X-ray diffraction (XRD) system, controlled by *CrysAlisPro* software (Rigaku Oxford Diffraction, 2022), following the protocol outlined in Atalay et al. (2022). The airflow temperature of Oxford Cryosystems’s Cryostream 800 Plus was set to 300 K (26.85 °C) and maintained consistently at ambient temperature during data collection. The 72-well Terasaki plate was mounted on the adapted XtalCheck-S plate reader attachment on the goniometer omega stage (Gul et al., 2023). Two dozen crystals were used for the initial screening to assess diffraction quality. Adjustments to omega and theta angles and X, Y, and Z coordinates were made to center crystals at the eucentric height of the X-ray focusing region. Subsequently, diffraction data were collected from multiple crystals. Crystals demonstrating the highest resolution Bragg diffraction peaks were chosen for further data collection, and exposure times were optimized accordingly to minimize radiation damage during overnight data collection. The most promising crystals were cultivated in a buffer solution containing 2.4 M NaCl, 100 mM Tris-HCl at pH 7.4, 6 mM ZnCI_2_, 20 (w/v) poly(ethylene glycol) PEG-8000. Throughout the data collection process, *XtalCheck-S* was configured to oscillate to the maximum extent permitted by the detector distance, aiming to optimize crystal exposure oscillation angles. 42 frames were captured, each running lasting 3 minutes (5 seconds per frame) for all individual crystals. Parameters were set with the detector distance at 100.00 mm, a scan width of 0.5 degrees oscillation, and an exposure time of 10.00 seconds per image.

#### Data processing and structure determination

Following the optimization of plate screening parameters for all crystals, a 42-degree data collection was conducted for each selected crystal. All crystals were organized in *CrysAlisPro* for comprehensive data collection. An optimal unit cell was selected, and peak identification and masking procedures were executed on the collected data. Subsequently, a batch script incorporating the `xx proffitbatch` command was generated for cumulative data collection. The batch data reduction was accomplished using the *CrysAlisPro* suite through the script command, producing files containing all integrated, unmerged, and unscaled data (*.rrpprof) for each dataset. To merge all datasets into a reflection data (.mtz) file, the *proffit merge* process in the *Data Reduction* section of the main window of *CrysAlisPro* was employed. The reduced datasets (.rrpprof files) were further merged using *proffit merge*. The entirety of the data underwent refinement, integration, and scaling processes through the aimless and pointless implementations within *CCP4* (Evans, 2014). Ultimately, the processed data were exported into *.mtz formats. The crystal structure of INSv was determined under ambient temperature within the R3:H space group, employing the automated molecular replacement tool *PHASER* integrated into the *PHENIX* software package (Adams et al., 2010). Utilizing a previously published X-ray structure (PDB ID: 4GBN) (Palmieri et al., 2013) as the initial search model, the coordinates from 4GBN were employed for the preliminary rigid-body refinement within *PHENIX*. Subsequent refinement steps included simulated-annealing refinement, adjustment of individual coordinates, and Translation/Libration/Screw (TLS) parameters. Furthermore, a refinement technique known as composite omit map, embedded in *PHENIX*, was executed to identify potential locations of altered side chains and water molecules. The final model underwent meticulous examination and model-building in *COOT*, focusing on positions exhibiting notable difference density. Extraneous water molecules located beyond regions of significant electron density were manually excluded. The X-ray crystal structure was visualized using *PyMOL* and *COOT* (Emsley et al., 2004).

## Results

### Novel double-substituted insulin variant (INSv) induces cellular activity

To enhance the flexibility of insulin, we introduced double substitution from arginine (Arg) to lysine (Lys) and proline (Pro) to aspartic acid (Asp) in chain B. The internal Arg22 on insulin was specifically mutated to Lys to mitigate the formation of undesired cleavage products, as trypsin exhibits a higher affinity for Arg than Lys. INSv was produced and subsequently folded through a refolding process in E. coli Rosetta™ 2 Competent Cells, which contained a double mutant INSv vector (Figure S1). Calcium imaging was performed to verify whether the INSv monomer induces any cellular activity compared to the IAsp monomer (Fig. 1). Calcium is an essential intracellular messenger in neurons; consequently, while most neurons exhibit a resting intracellular calcium concentration ranging from approximately 50 to 100 nM, this concentration may transiently escalate by a factor of 10 to 100 during electrical activity (Berridge et al., 2000). Intracellular calcium signals operate a broad spectrum of processes, from neurotransmitter release on a microsecond scale to gene transcription occurring over minutes to hours. In examining spontaneous activity, we measure the number of events, which are the occurrences noted at each Region of Interest (ROI). Another parameter we analyze is the Area Under Curve (AUC), highlighted in red on the plot of spontaneous activity (Figure S2). Upon the individual administration of INSv vs. IAsp monomers to the cells, the count of cells demonstrating calcium activity increased, accompanied by a decrease observed in the Area Under the Curve (AUC) (Fig. 1C; Figure S2-S3; Movie S1, and Movie S2). The INSv and IAsp monomers initiated the neural networking, resulting in a decrease in AUC, which is expected as the event digit decreases. However, while the AUC decrease in IAsp is deemed insignificant, that observed in INSv is considered significant.

**Figure 1.**
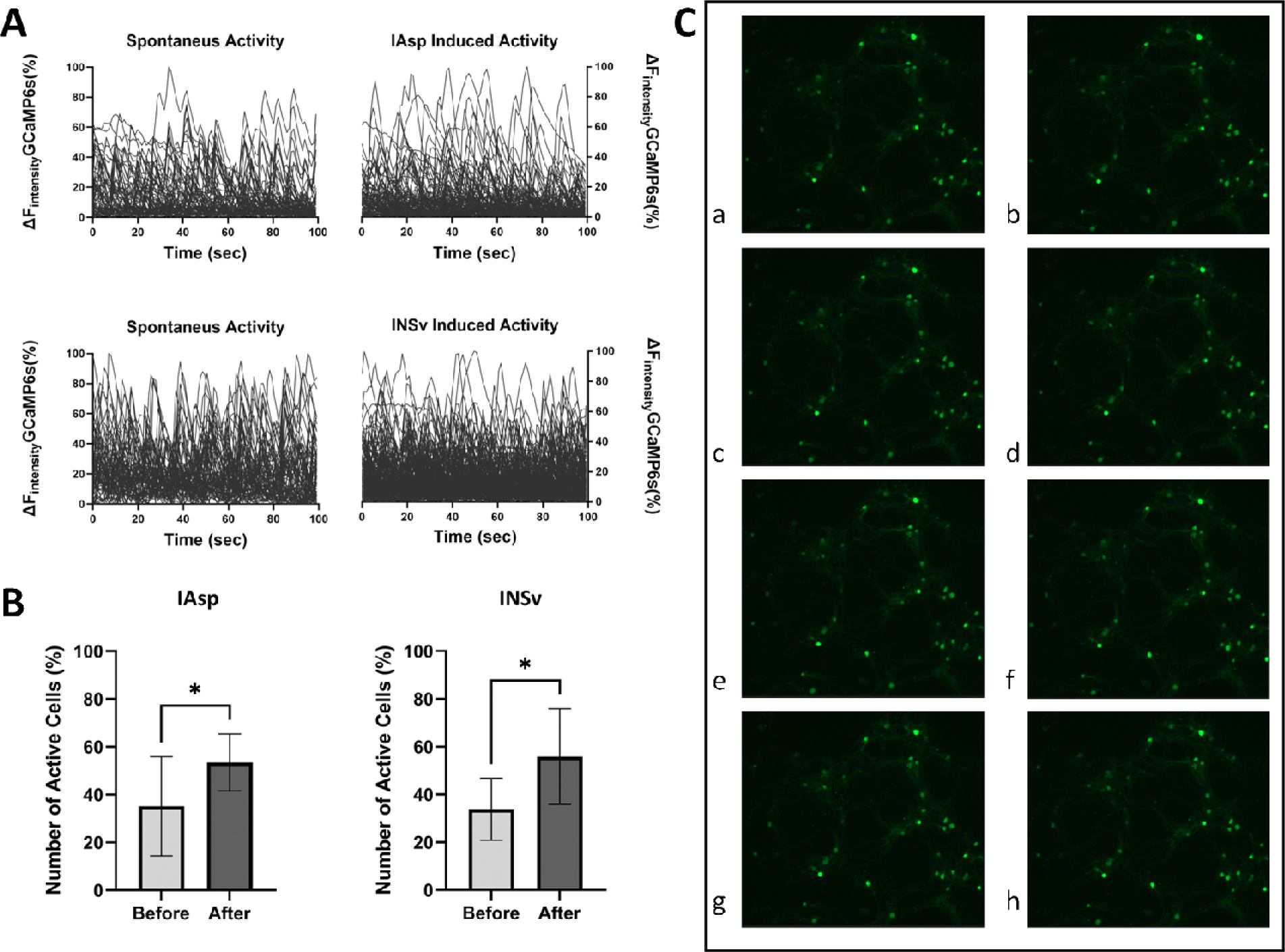
Comparative analysis of calcium imaging between INSv and IAsp. **(A)** The graph display temporal light intensity shifts indicative of recorded calcium activity. **(B)** Statistical analysis of the effect of INSv and IAsp on the proportion of active neural cells was performed using an unpaired t-test, yielding a p-value of 0.0358 for INSv and 0.0473 for IAsp. **(C)** Sequential visualization of calcium activity propagation following INSv stimulation includes **a)** a pre-stimulation network image and **b-h)** a progressive spread of calcium activity across the network, with detailed annotations for each stage.

### Alternate dynamic status in IAsp and INSv monomers

In this study, we quantified deuterium exchange in distinct regions of INSv and IAsp, identified the locations of disparities, and elucidated the variations in their local dynamics. In conducting HDX-MS reactions, we specifically opted for the monomeric form of insulins to determine differences in monomer dynamics under identical experimental conditions. Consequently, even if substantial dynamic differences were not observed in the mutated region of these insulins, notable differences were noted in the dynamics of the remaining monomeric regions (Fig. 2 and Fig. 3).

**Figure 2.**
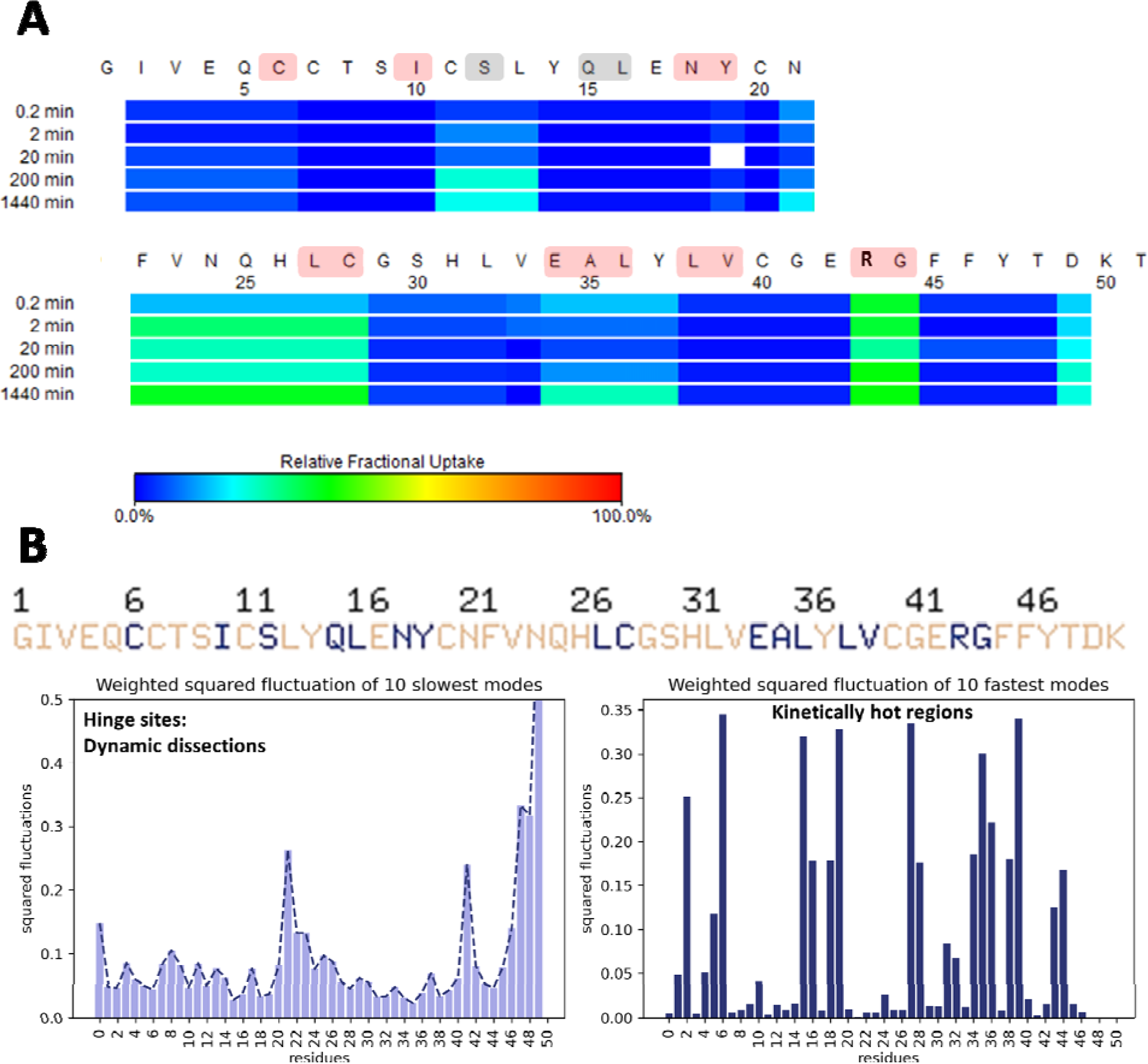
Complementary experimental and computational dynamic analysis of IAsp. **(A)** Heat map displaying time-course H/D exchange measurements of IAsp. The color code from blue to red describes deuterium exchange. **(B)** Cumulative 10 slowest and 10 fastest modes from GNM to show dynamic dissections and kinetically hot regions. The critical residues for relative fractional deuterium exchange, which are highlighted in red, are largely consistent with the cumulative 10 slowest and 10 fastest modes from GNM. The residues observed in the H/D reaction alone are illuminated in gray.

**Figure 3.**
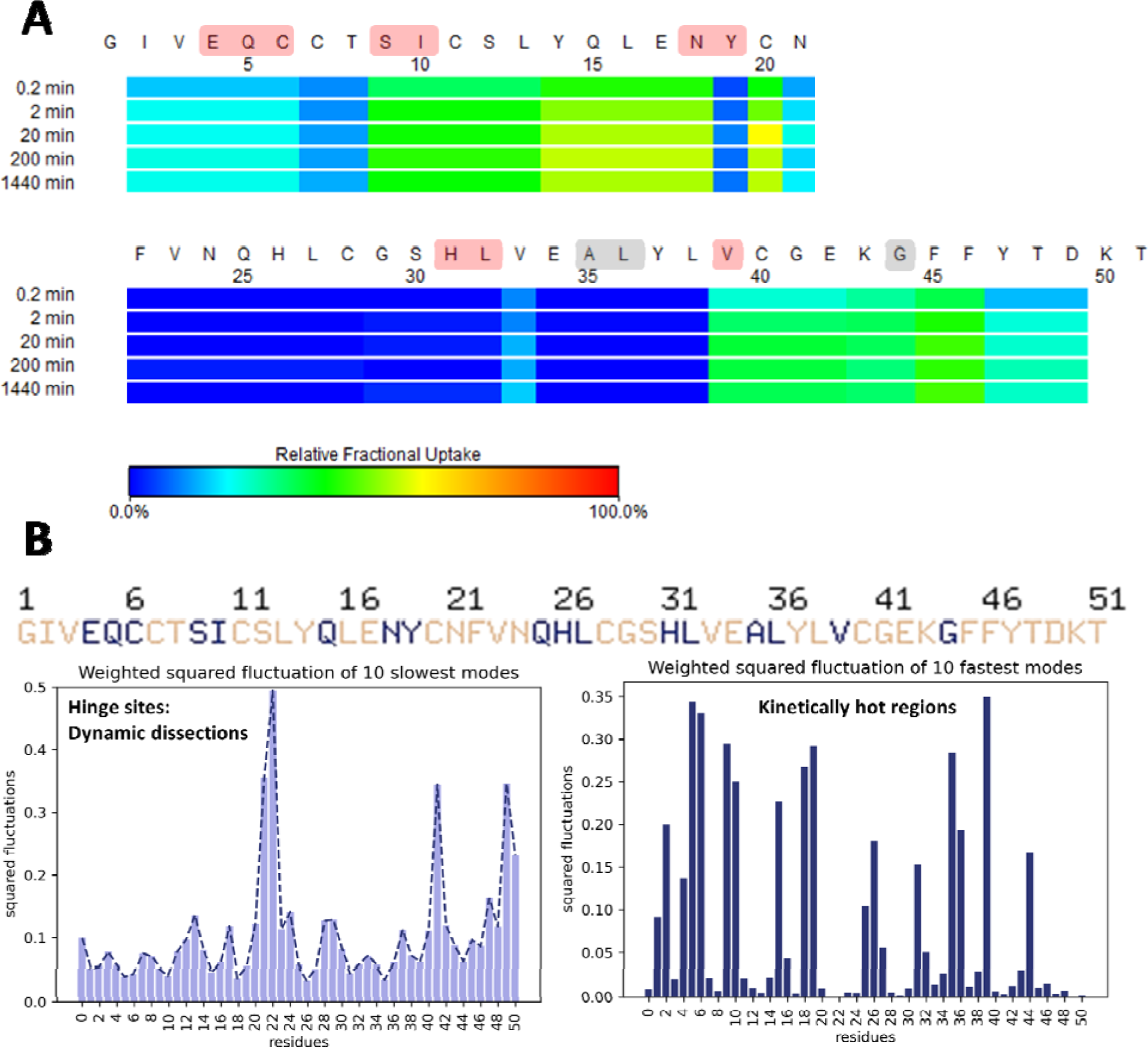
Complementary experimental and computational dynamic analysis of INSv. **(A)** Heat map displaying time-course H/D exchange measurements of INSv. **(B)** Cumulative 10 slowest and 10 fastest modes from GNM to show dynamic dissections and kinetically hot regions. The critical residues for relative fractional deuterium exchange, which are highlighted in red, are largely consistent with the cumulative 10 slowest and 10 fastest modes from GNM. The residues observed in the HDX reaction alone are illuminated in gray in panel A.

Initially, to ensure uniformity in pH conditions during the exchange reaction, Tris-HCl buffer at pH 9.1 (pD 9.5) was selected, owing to its ability to maintain insulin’s *in vivo* monomeric state (REF). The temporal evolution of deuterium exchange profiles by IAsp is indicated in Figure 2. In the H/D exchange reaction of the IAsp monomer, certain residues, namely regions involving Cys11-Leu13, Asn21-Cys28, Glu34-Tyr37, Arg43-Gly44, and Asp49, exhibit a notable degree of deuterium exchange indicating relatively more mobile compared to other residues (Fig. 2A), consistent with findings from computational analyses (Fig. 2B). Our normal mode analysis identifies Ile2, Cys6, Ile10, Gln15-Leu16, Asn18-Tyr19, Leu27-Cys28, Glu34-Leu36, Leu38-Val39, and Arg43-Gly44 as regions undergoing dynamic dissections, and these are referred to as kinetically hot regions (Fig. 2B). These identified hinge sites from our computational analysis corroborates with observations in the HDX-MS heat-map results. However, Ser12 and Gln15-Leu16 residues are identified as crucial hinge regions in only GNM analysis (Fig. 2B).

Next, the H/D exchange reaction in INSv insulin analog was compared to IAsp to assess potential distinctions. The INSv analog demonstrated higher deuterium exchange compared to IAsp (after a 1440-minute reaction, n = 3), displaying a more dynamic conformation for INSv analog compared to IAsp, agreeing with the findings of DSC analysis (Fig. 3A). Within the H/D exchange reaction of the INSv monomer, specific regions, Ile2-Cys6, Ser9-Asn18, Cys20-Asn21, and Val39-Asp49, exhibited a noticeable degree of relative mobility in contrast to other residues, broadly consistent with computational analyses (Fig. 3A). Our normal mode analysis identified Glu4-Cys6, Ser9-Ile10, Asn18-Tyr19, His31-Leu32, and Val39 residues as regions undergoing dynamic dissections, denoted as kinetically hot regions (Fig. 3B). Notably, the identified hinge sites from our computational analysis align seamlessly with observations in th HDX-MS heat-map results. Nevertheless, Ala35-Leu36 and Gly44 residues consistently emerged as crucial hinge regions only in our GNM analysis (Fig. 3B).

Figure 4A depicts the close-up view of deuteration levels of the corresponding fragments of INSv and IAsp over the 200-minute time course. The uptake profiles of most fragments exhibited variations between INSv and IAsp (Fig. 4A-B, Figure S4 and Figure S5). Additionally, there was a minimal resemblance in the fragment encompassing His31-Asp49, which includes the mutation site in INSv (Fig. 4B).

**Figure 4.**
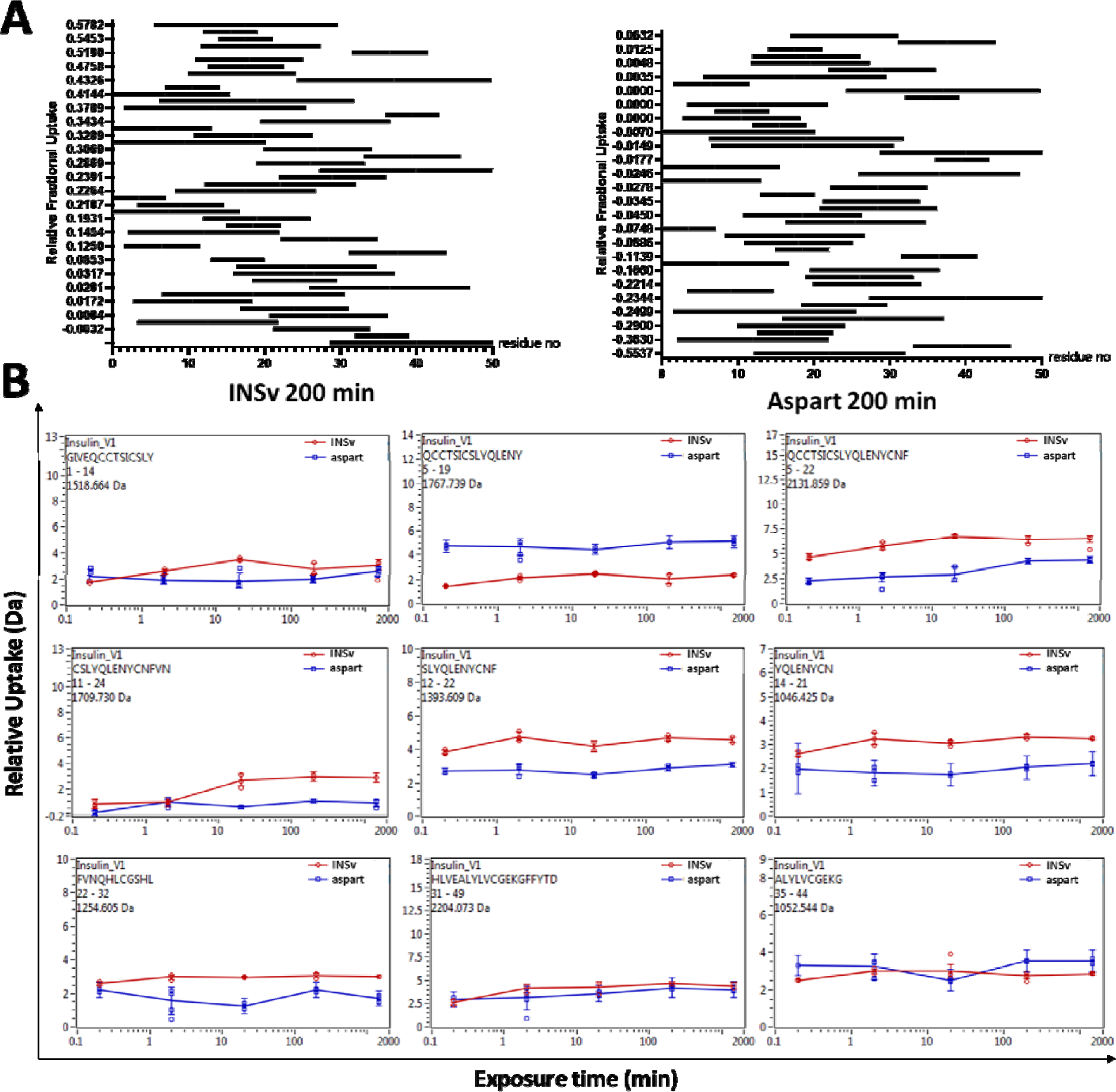
Projection of the peptide-specific deuterium exchange. **(A)** Relative deuterium exchange of INSv an IAsp peptides depicted by Woods plots over the 200 min time course. **(B)** Comparison between time-dependent relative deuterium exchange of representative INSv and IAsp peptides. Data is represented on a logarithmic scale.

### Thermal stability analysis confirms distinctive H/D dynamics of both conformers

Differential scanning calorimetry (DSC) measurements were conducted to investigate the thermal stability of the INSv compared to commercially available IAsp. DSC measurements revealed significant differences in the thermodynamic properties of IAsp and INSv (Fig. 5). The thermal unfolding of IAsp shows a steep endothermic denaturation with a T_m_ of 81.9 °C, a ΔH_cal_ of 35.46 KJ/mol, and a ΔS of 0.099 KJ/mol·K. On the other hand, the T_m_ of the INSv was considerably lower (T_m_ = 77.6 °C), and the ΔH_cal_ also decreased (ΔH_cal_= 31.55 KJ/mol), as well as the ΔS showed increasing tendency (ΔS= 0.142 KJ/mol·K). The significantly lower T_m_ suggests a less stable and, most likely, less compact conformation for the INSv. Moreover, the decreased ΔH_cal_ and increased ΔS of INSv align with the lower T_m,_ assuming the protein conformation is more flexible (Fig. 5A). This interpretation is reinforced by the distinct H/D reactions observed between INSv and IAsp, indicating comparable flexibility of INSv relative to IAsp in H/D exchange (Fig. 5B).

**Figure 5.**
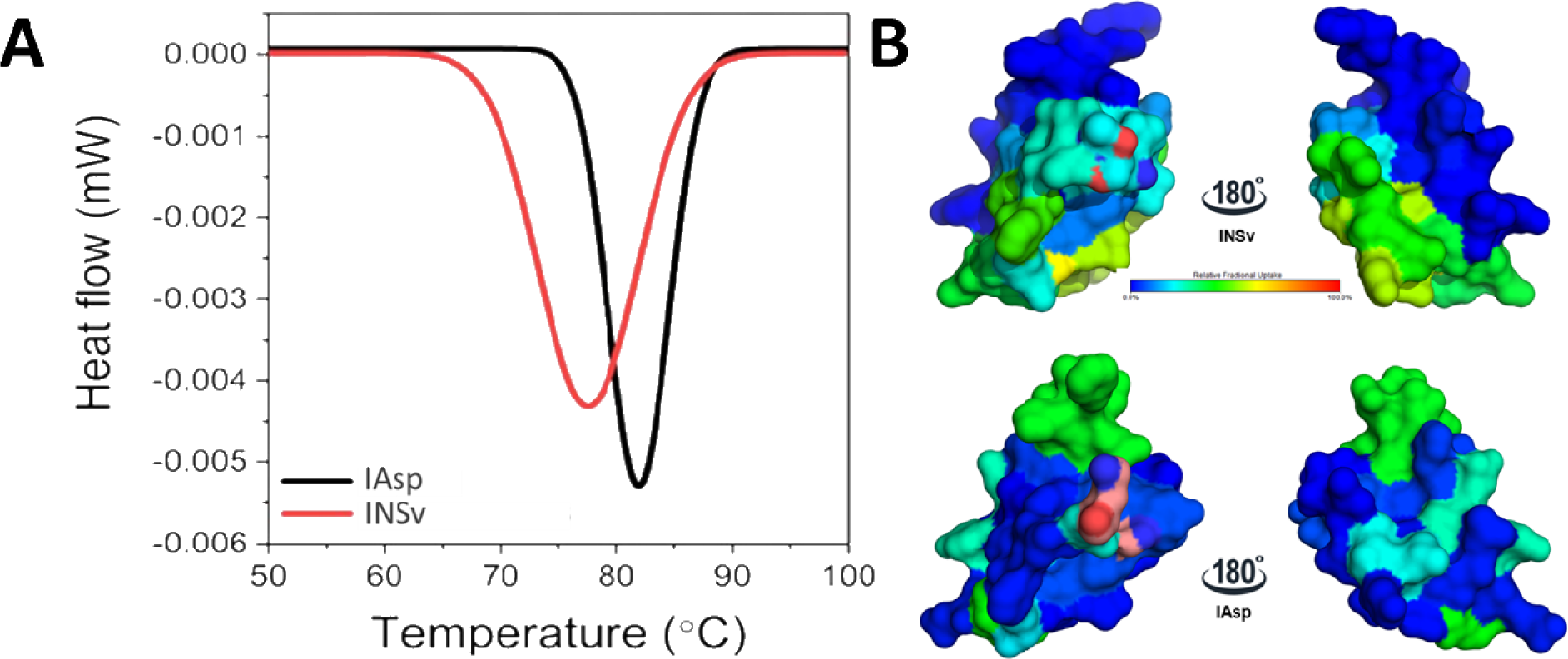
Representation of thermal stability analysis comparing to H/D dynamics variability of IAsp and INSv analogs. **(A)** The variation in heat flow depends on the melting temperature (Tm), which signifies the point at which 50% of the protein undergoes denaturation. Furthermore, the calorimetric enthalpy (ΔHcal) was computed, and the alteration in conformational entropy (ΔS) was ascertained by evaluating the ratio of ΔHcal to Tm in Kelvin (K). **(B)** Visualization of deuterium exchange over 1440-minute intervals through aligning our newly determined monomer of INSv with the monomer of IAsp. The blue spectrum marks the areas of rigidity, whereas the red spectrum indicates regions of flexibility. Figures have been generated by *PyMOL.*

### Crystal structure of hexamer INSv crystal form is compatible with in-solution analyses

We determined the ambient temperature structure of INSv at 2.5 Å resolution (Table 1 and Figure S6). Two nearly identical monomer molecules within the asymmetric unit cell exhibit an overall RMSD of 0.81 Å. Superpositioning and comparative analysis with the PDB structure of 4GBN revealed overall RMSD values ranging from 0.3 Å to 0.83 Å for the nearly identical monomer molecules in the asymmetric unit. Molecular replacement with a dimer in the asymmetric unit facilitated the solution of the crystal structures of INSv insulin at pH 7.5. Specifically, amino acid residues 1 to 8 exhibited a “T” conformation in one monomer and an “R” conformation in the other with well-defined electron densities (Fig. 6). In the previously reported IAsp structure (PDB_ID: 4GBN), three symmetry-related “TR” dimers constituted the biological unit (Palmieri et al., 2013). The hexameric configuration of INSv is organized near a crystallographic 3-fold axis aligned with the central channel (Palmieri et al., 2013). In contrast to the 4GBN structure, a distinctive feature of the INSv hexamer involves the outward orientation of Asn3 in chain B relative to the interior of the channel (Fig 6A-B). This spatial arrangement produces less stable electrostatic interaction with the two symmetry-related B3Asn residues (Fig. 6). The crystal structures of INSv and 4GBN exhibit similarities in both Cα atoms and side chains. Although there are positional variations, these are primarily minor, with relatively major discrepancies observed in the side chains of 4GBN (Palmieri et al., 2013) IAsp compared to INSv. Upon superimposing the current INSv insulin structure onto 4GBN in the T_3_R_3_ conformation, a dissimilarity becomes evident in the region encompassing amino acid residues 1 to 8 in chain B. This dissimilarity is observed in both backbone and side chain conformations, providing evidence for INSv insulin adopting the T_3_R_3_ conformation with a pure TR-state dimer configuration. A pairwise global structural alignment between the T_3_R_3_-INSv and -4GBN IAsp structures highlights their similarity, with a global RMSD slightly over 0.8 Å. This similarity persists, indicating a consistent outcome attributable to the zinc-bound form.

**Figure 6.**
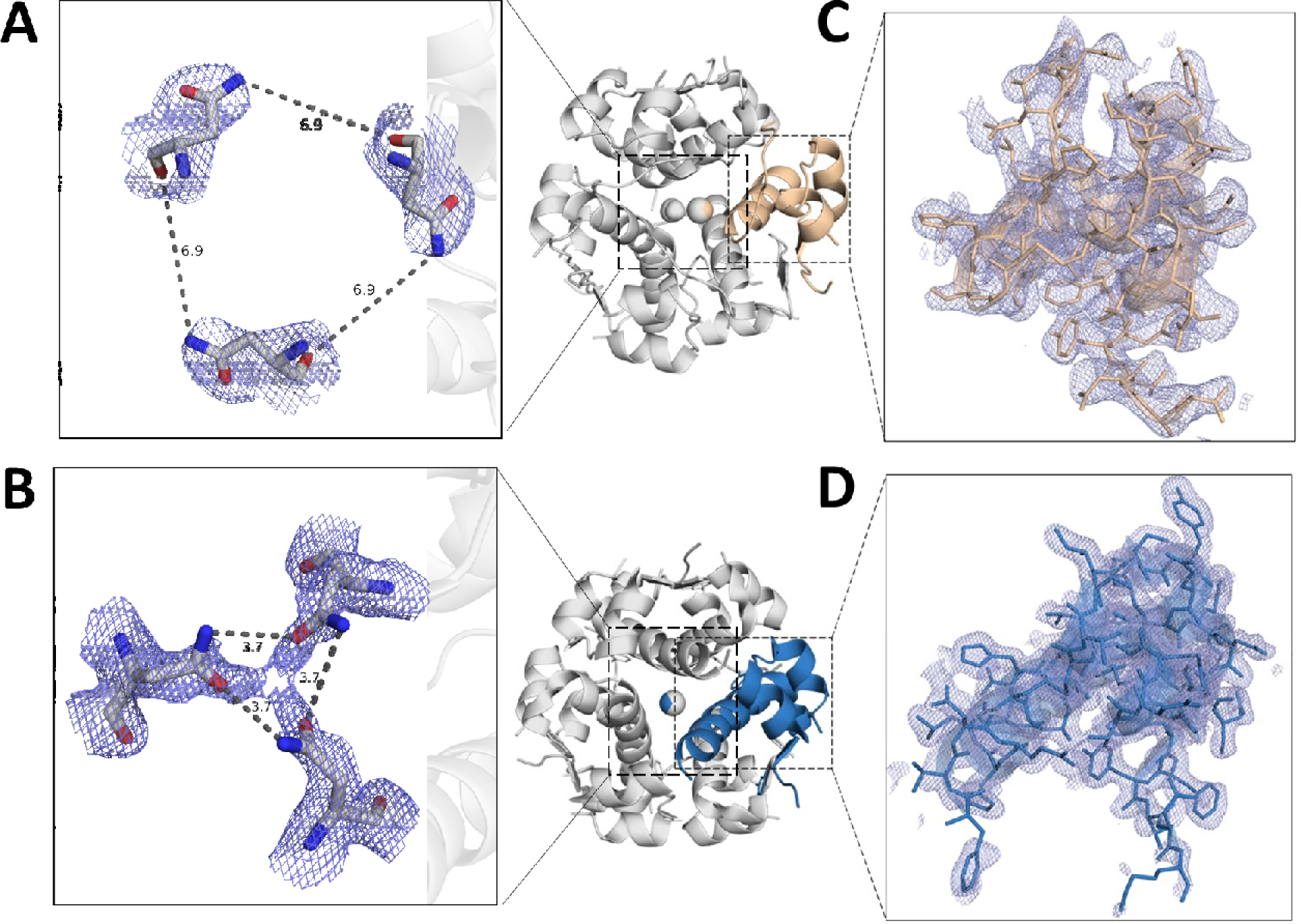
A comparison between the INSv and 4GBN IAsp structures, presenting both distant and close-up views. Panels **(A)** and **(B)** depict the electrostatic interaction distances of AsnB3 residues within each monomer of insulin hexamers, INSv and 4GBN, respectively. Additionally, panels **(C)** and **(D)** examine a monomer from insulin hexamers INSv and 4GBN, respectively. 2*Fo-Fc* simulated annealing-omit map at 1 sigma level is colored in slate.

**Table 1.** Data collection and refinement statistics of *INSv* insulin analog.

|  |  |
| --- | --- |
| **Data collection** | *INSv* |
| **Instrument** | *Turkish DeLight* |
| **Resolution range** | 21.64 - 2.501 (2.59 - 2.501) |
| **Space group** | R 3: H |
| **Unit cell** | 80.376 80.376 38.04 90 90 120 |
| **Redundancy** | 24.9 (12.9) |
| **Completeness (%)** | 97.70 (91.02) |
| **Mean I/sigma(I)** | 6.88 (1.72) |
| **Wilson B-factor** | 37.28 |
| **R-merge** | 0.6092 (1.776) |
| **R-meas** | 0.6197 (1.849) |
| **R-pim** | 0.1113 (0.502) |
| **CC1/2** | 0.952 (0.358) |
| **CC\*** | 0.988 (0.726) |
| **R-work** | 0.3069 (0.4477) |
| **R-free** | 0.3529 (0.3590) |
| **CC (work)** | 0.846 (0.192) |
| **CC (free)** | 0.689 (0.600) |
| **Protein residues** | 102 |
| **RMS (bonds)** | 0.091 |
| **RMS (angles)** | 0.86 |
| **Ramachandran favored (%)** | 94.68 |
| **Ramachandran allowed (%)** | 4.26 |
| **Ramachandran outliers (%)** | 1.06 |
| **Average B-factor** | 55.83 |
| **Macromolecules** | 56.44 |
| **Ligands** | 38.47 |
| **Solvent** | 45.06 |

We conducted a separate B-factor analysis for the monomers of 4GBN and INSv (Fig. 7). In the R-forms of the INSv and 4GBN IAsp structures, it is observed that the N-terminal segments ranging from Asn21-Gln25 and Gly41-Glu42 of INSv’s B chain exhibit slightly higher fluctuations compared to their counterparts in the 4GBN IAsp structure. Similarly, in the T-forms of the INSv and 4GBN IAsp structures, the regions spanning Cys40-Glu42 in INSv demonstrate pronounced flexibility relative to the T-form of the 4GBN IAsp structure. On the other hand, the R-form of INSv is greatly compatible with our H/D reactions pattern we observed in Figure 3, assuming INSv monomer in-solution is probably a more flexible R-form, thus implying that the R-forms of INSv monomer exhibit greater fluctuation than its T-form. Additionally, substituting R with K might induce a flexible R-state despite the limited influence of Zn^+2^ coordination on the hexameric form.

**Figure 7.**
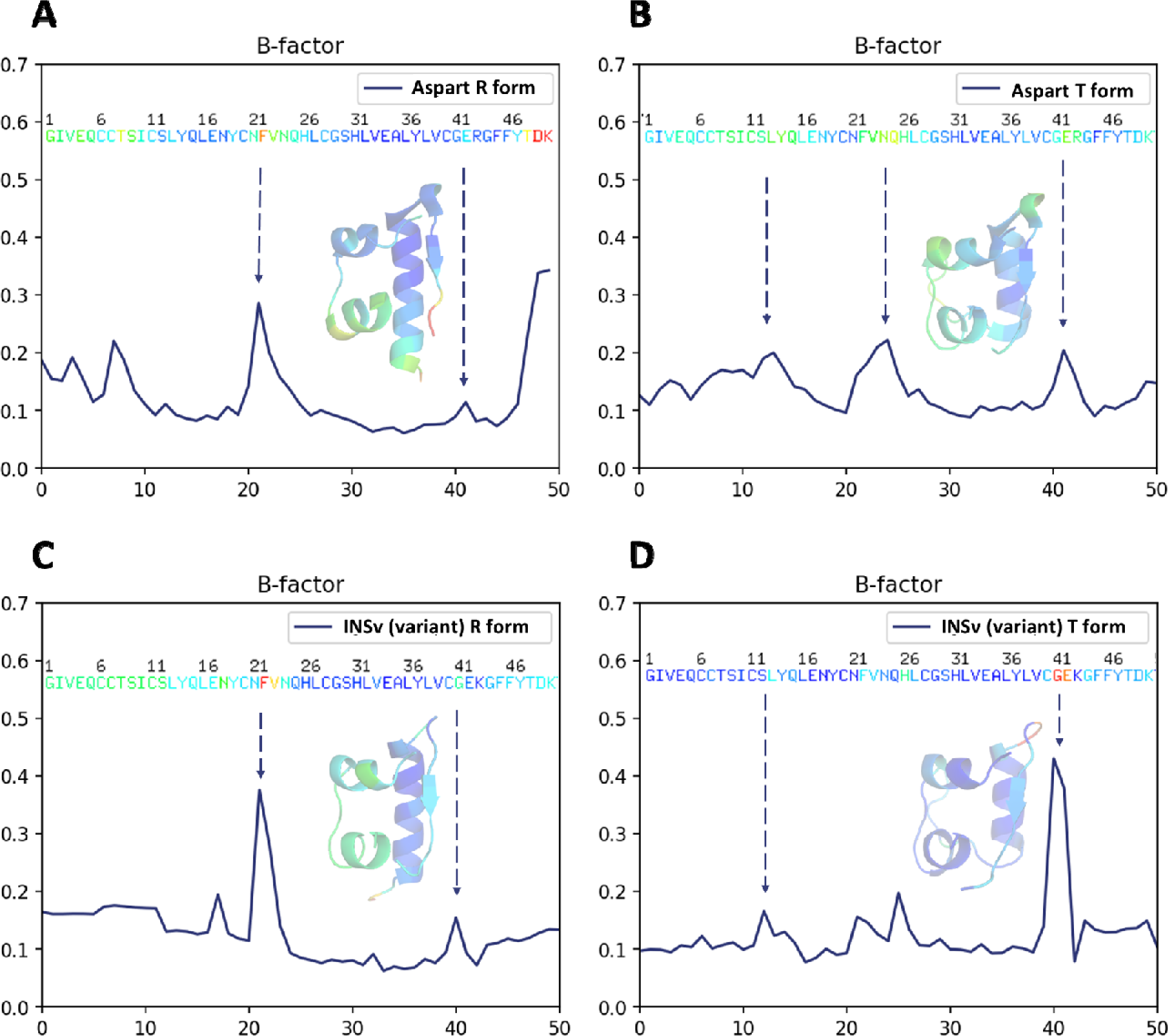
Experimental B-factor analysis and cartoon representation of each 4GB IAsp and INSv dimers’ monomer. **(A, B)** Thermal distance analysis supports the experimental b-factor of the IAsp structure’s R-state and T-state conformers. **(C, D)** Thermal distance analysis supports the experimental b-factor of the INSv structure’s R state an T state conformer. Cartoon mode is colored by the B-factor in PyMOL. The blue spectrum indicates areas of rigidity, whereas the red spectrum indicates regions of flexibility.

## Discussion

The zinc^+2^-coordinated hexameric form of insulins exemplifies the characteristic structural model inherent in globular proteins (Chothia et al., 1983). Serving as a pivotal model in the evolution and application of X-ray crystallographic techniques (Baker et al., 1988), the hexamer functions as a stable reservoir of the hormone within both the secretory granules of pancreatic β-cells and pharmaceutical formulations (Brange et al., 2012). The integration of structure-based mutagenesis into the Zn^+2^-bound insulin hexamer (Brange et al., 2012; Dodson and Steiner, 1998) has significantly advanced the development of rapid-acting insulin analog formulations suitable for prandial injection and pump use (Zinman et al., 1997). Despite their initial categorization as “monomeric” (Brange et al., 2012), these analogs are presently designed either as Zn^+2^-bound insulin hexamers or as an equilibrium distribution of Zn^+2^-free oligomers (DeFelippis et al., 2001; Garg et al., 2005). Although it assembles into dimers and hexamers during its biosynthesis and storage, insulin is always active as a monomer (Ayan, 2022). Even though X-ray analysis can reveal the 3-dimensional structure of the insulin monomer states (Nagata et al., 1995; Kurapkat et al., 1999), studies about the monomer structure and dynamics in solution are severely limited by insulin self-association into dimers and higher oligomers. In a study about two main conformations of insulin which differ in the extent of the helix in the B chain (B9-B20 and B1-B20, respectively) have been identified, demonstrating that reagents such as chloride and phenol govern the conformations present in the insulin hexamers and can influence the behavior and properties of insulin preparations employing them (Derewenda et al., 1989). In a study about to be used nuclear magnetic resonance (NMR) spectroscopy, they show that the monomer insulin is generally similar to crystal structures and resembles a crystalline T-state more than an R-state in the sense that the B-chain helix is confined to residues B9-B19 (Olsen et al., 1996). In an NMR-derived human insulin monomer structure in water/acetonitrile study, they used two different molecular dynamics simulated annealing refinement protocols. Observing that the structures of monomer that form part of a hexamer in the X-ray structure and in water/acetonitrile solution overlay very well both in terms of backbone and side chains (Bocian et al., 2008).

Assorted experimental evidence suggests that monomeric insulin exhibits significant conformational heterogeneity, and modifications of apparently disordered regions affect both biological activity and the longevity of pharmaceutical formulations, presumably through receptor binding and fibrillation/degradation, respectively (Busto-Moner et al., 2021). The present study has explored the examination of the functional (Fig. 1 and Figure S2-S3) INSv alongside the IAsp monomers to illuminate the fundamental structure and dynamics of these rapid-acting insulin in a solution. The substitution of Arg with Lys at position B22 and Pro with Asp at position B28 distinguish the INSv, which DSC confirmed to be a less stable insulin analog compared to IAsp. Lys and Arg, both positively charged basic amino acids under physiological conditions, are predominantly exposed on the surfaces of proteins (Galligan et al., 2017; Li et al., 2013; Warwicker et al., 2014). Despite their shared function as basic residues, Arg contributes more stability to protein structure than Lys, attributed to its distinctive geometric structure (Sokalingam et al., 2012). The guanidinium group in Arg facilitates interactions in three possible directions over its three asymmetrical nitrogen atoms (Nη1, Nη2, Nε). In contrast, the basic functional group of Lys allows interaction in only one direction (Xu et al., 2017; Malinska et al., 2016; Wu et al., 2016). This capability enables Arg to form more electrostatic interactions, including salt bridges and hydrogen bonds, than Lys. Presumably, this results in stronger interactions than those generated by Lys.

Based on the compatibility between the experimental B-factor analysis and the H/D exchange reaction of INSv (Fig. 3 and Fig. 7), the in-solution structure of this variant (Fig. 3) closely mirrors the crystallographic R state (Fig. 7). However, unlike the IAsp structure, a distinct feature of the INSv hexamer in the R form involves the outward orientation of Asn3 in chain B relative to the interior of the channel (Fig. 6). This spatial configuration leads to a less stable electrostatic interaction with the two symmetry-related AsnB3 residues. Additionally, the unique R-state of the INSv crystal structure is evident in a flexible INSv monomer according to B-factor analysis (Fig. 7), which correlates with the HDX-MS findings for the INSv monomer (Fig. 3). Characterizing INSv and IAsp as native-like monomers at pH 9.0 (Tokumoto et al., 2006; Fredericq, 1953; Kadima et al., 1992) and without an organic co-solvent provides an opportunity for a quantitative assessment of amide-proton exchange in D_2_O. HDX-MS data for INSv indicated swift deuterium exchange at the initial time points in the majority of peptides, followed by local exchange after specific time points (Fig. 3A). In contrast, IAsp exhibited no deuterium exchange in the first 35 residues, with only the Phe22-Cys28 and Arg43-Gly44 peptides showing deuterium exchange and local exchange at 2-min, 20-min, 200-min and 1440-min points (Fig. 2A).

The formation of insulin oligomers primarily relies on establishing noncovalent bonds between interacting residues from adjacent monomers (Dunn, 2005). However, alterations in pH can interrupt these interactions, causing residues in distant sites to cease their communication paths and transition into monomeric states (Fredericq, 1953; Tokumoto et al., 2006). Modifying monomeric states results in substantial modifications to the customary residue communication network compared to oligomeric states (Bolli et al., 2000; Eren et al., 2021). These alterations can also be elucidated through computational GNM analysis (Bakan et al.,2011), which is analogous to our insights from experimental H/D exchange reactions (Fig. 2B and Fig. 3B). We conducted GNM in this study to further elucidate the distinctive dynamics exhibited by the INSv monomer compared to the monomeric form of IAsp, aiming to correlate HDX-MS results. The monomeric IAsp (NovoRapid^TM^) was employed as the reference model, which is analogous to the methodology used in HDX-MS. GNM-based methodologies precisely assess global protein motions at lower frequencies and local motions at higher frequencies (Ayan et al., 2022; Eren et al., 2021). In the INSv monomer, pivotal hinge sites crucial for insulin activity are identified in residues Cys6, Ser9, Asn18-Tyr19, and Leu32, as observed through GNM and HDX-MS analyses (Fig. 3B). Conversely, in the IAsp monomer, hinge sites critical for insulin activity are in Cys6, Ile10, Asn18-Tyr19, Cys28, Glu34, Leu38, and Arg43-Gly44 (Fig. 2B). Remarkably, these residues (in Fig. 2B and Fig. 3B) exhibiting dynamic changes are the regions involved in receptor interactions (De Meyts, 2000).

Finally, observed that both INSv (before, 34% activity; after, 56% activity) and IAsp (before, 35% activity; after, 53% activity) significantly augment the activity of cells (Fig.1; Figure S2-S3), but they do not differ considerably from each other in the calcium imaging experiment. In both cases, there is an increase in the active cell count due to spontaneous events. Based on a study that used Fura-2 instead of GcaMP6, it was claimed that Fura-2 might not be sufficient to detect all events as well as suggested that the insulin concentration should be increased (Maimaiti et al., 2017). Accordingly, our experiment was designed with a relatively higher insulin dosage of 3.4 µM and utilized GcaMP6 instead of Fura-2. Although GcaMP6 and Fura-2 are highly similar, GcaMP6 is more effective at detecting smaller events (Garcia et al., 2017). Quantitative analysis shows that the pattern of spontaneous activity in INSv is denser than in IAsp, likely due to distinct calcium transients. This indicates that the calcium influx, indicative of the electrical activity of the cells, is more significant in INSv than in IAsp. Additionally, both analogs acutely induce neural networking, accompanied by a decline in AUC values. The reduction in AUC for IAsp is insignificant (*p*-value: 0.1290), while the decrease in AUC for INSv is statistically significant (*p*-value: 0.0002) (Figure S2). This indicates that the observed disparity between INSv and IAsp is due to the higher number of events in INSv compared to IAsp. This suggests that the distinct underlying mechanisms may be related to the different kinetics of INSv and IAsp, as observed *in solution* and *in crystallo* experiments (Fig. 2; Fig. 3; Fig. 5 and Fig. 6). The dynamic variations observed, both experimentally and computationally, suggest that while the hexameric structure bound to the Zn^+2^ cation may not change significantly, the substitution of Arg with Lys in a monomer induces distinct kinetic models dependent on particular mobilities. Considering the tendency of conformational fluctuations to hasten the degradation of pharmaceutical formulations, we propose that adopting a concept referred to as ’dynamic restructuring’ for insulin could enable the strategic development of formulations with smart rapid-acting, given the observed kinetic differences.

## Conclusion

In this study, we introduced a less stable & functional monomeric insulin variant, INSv. The crystal structure of its hexameric insulin variant was determined at a resolution of 2.5 Å at ambient temperature. A H/D exchange dynamic between the IAsp and the INSv revealed that INSv exchanges deuterium atoms more rapidly and extensively than intact IAsp, which is consistent with the diminished hexameric stability of INSv. GNM analysis further supported these kinetic differences. The observed variations prompted exploring the specific regions responsible for both molecules’ discrepant H/D exchange reactivity. These deuteration discrepancies, coupled with computational analysis, imply that the R→K substitution can induce distinct kinetic models, even under identical physiological conditions.

## Ethics approval and consent to participate

Not applicable.

## Consent for publication

Not applicable.

## Availability of data and materials

INSv (LysB22-AspB28) analog has been submitted to PDB under the 8Z4B accession code.

## Authors’ contributions

E.A., M.I.A., H.D., and G.D.D. designed the experiments. E.A. prepared the samples. E.A. performed the sample delivery, data collection, and data processing. Structures were refined by E.A. DSC analysis was performed by E.T. In-vitro experiment, and calcium imaging was performed by S.B., E.A., and M.I.A. HDX-MS analysis was performed by M.T., O.T., E.A., and B.D. E.A performed GNM analysis. E.A., M.T., O.T., E.T., B.D., S.B., M.I.A., G.D.D., and H.D. prepared the manuscript.

## Acknowledgment

The authors thank Dr. İrfan Çinkaya for his invaluable support. The authors gratefully acknowledge the use of the services and facilities of the Istanbul Technical University MOBGAM (Molecular Biology-Biotechnology&Genetics Research Center), University of Health Science-Validebag DETAUM (Experimental Medicine Research and Application Center), Istanbul Medipol University, Research Institute for Health Sciences and Technologies (SABITA) and DEVA Holding A.Ş.

---

**Supplementary Figure 1.**
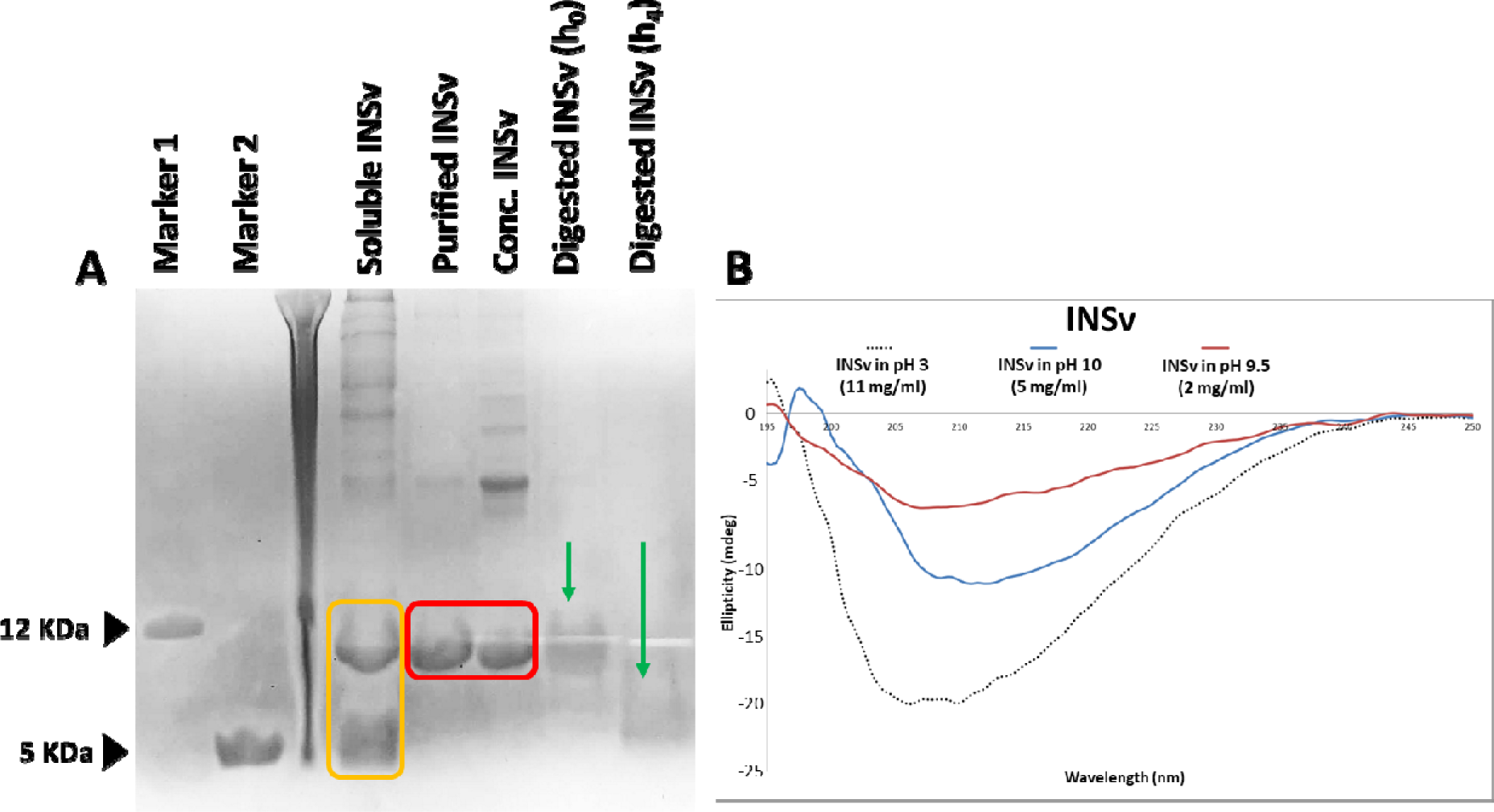
The monomeric INSv that produced by the refolding process. **(A)** 20% SDS-PAGE analysis of INSv, Marker 1 indicates proinsulin (Iodinated Human proinsulin, Sigma-Aldrich, 9015) (∼12 KDa), and Marker 2 indicates monomeric insulin aspart (NovoRapidTM, Novo Nordisk, Denmark) (∼5 kDa). Soluble INSv contains non-purified proINSv (12 KDa) and digested INSv (5 kDa), which is colored yellow. Purified and concentrated proINSv have been highlighted in red. INSv is performed enzymatic digestion with trypsin through zero time (h0) and 4-hour (h4) at 30°C, which are indicated by green arrows. Each line consists of 2.5 mg/ml pro- and mature INSv samples. **(B)** Circular Dichroism plot of distinct buffered monomeric INSv at pH 3, pH 10, and pH 9.5.

**Supplementary Figure 2.**
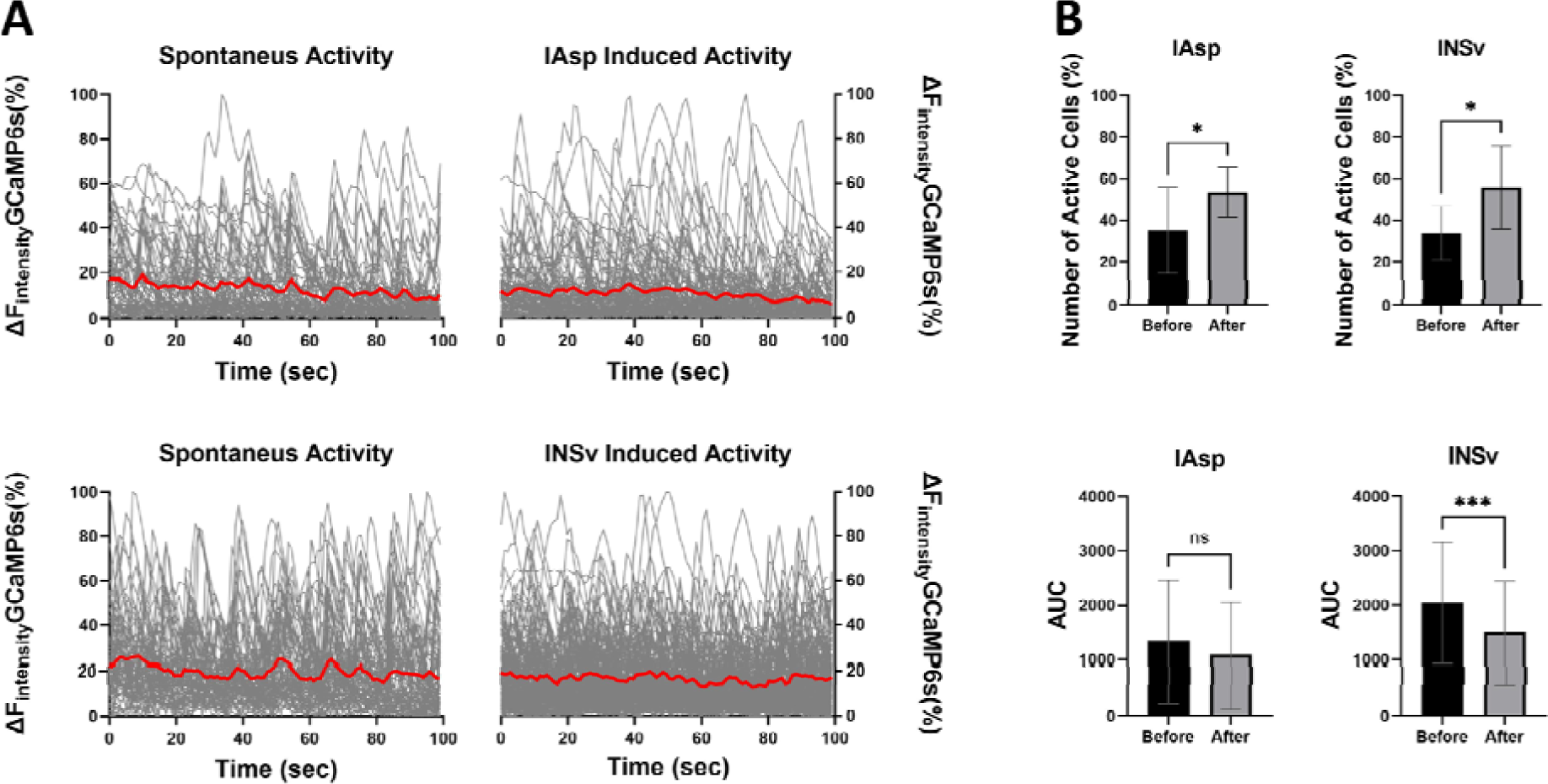
Comparative spontaneous activity analysis between INSv and IAsp. **(A)** The graph displays temporal light intensity shifts indicative of recorded calcium activity. The under areas of the red trend have highlighted the area under the curve (AUC), which indicates the average of the areas under the light intensity change curve recorded from each ROI. **(B)** Statistical analysis of the AUC measurement (left) indicated that INSv application significantly changed spontaneous calcium activity (IAsp *p*-value: 0.1290, INSv *p*-value: 0.0002). The mean AUC decreased in both samples. Statistical analysis of the effects of INSv and IAsp on the proportion of active neural cells (right) was performed using an unpaired t-test, yielding a *p*-value of 0.0358 for INSv and 0.0473 for IAsp.

**Supplementary Figure 3.**
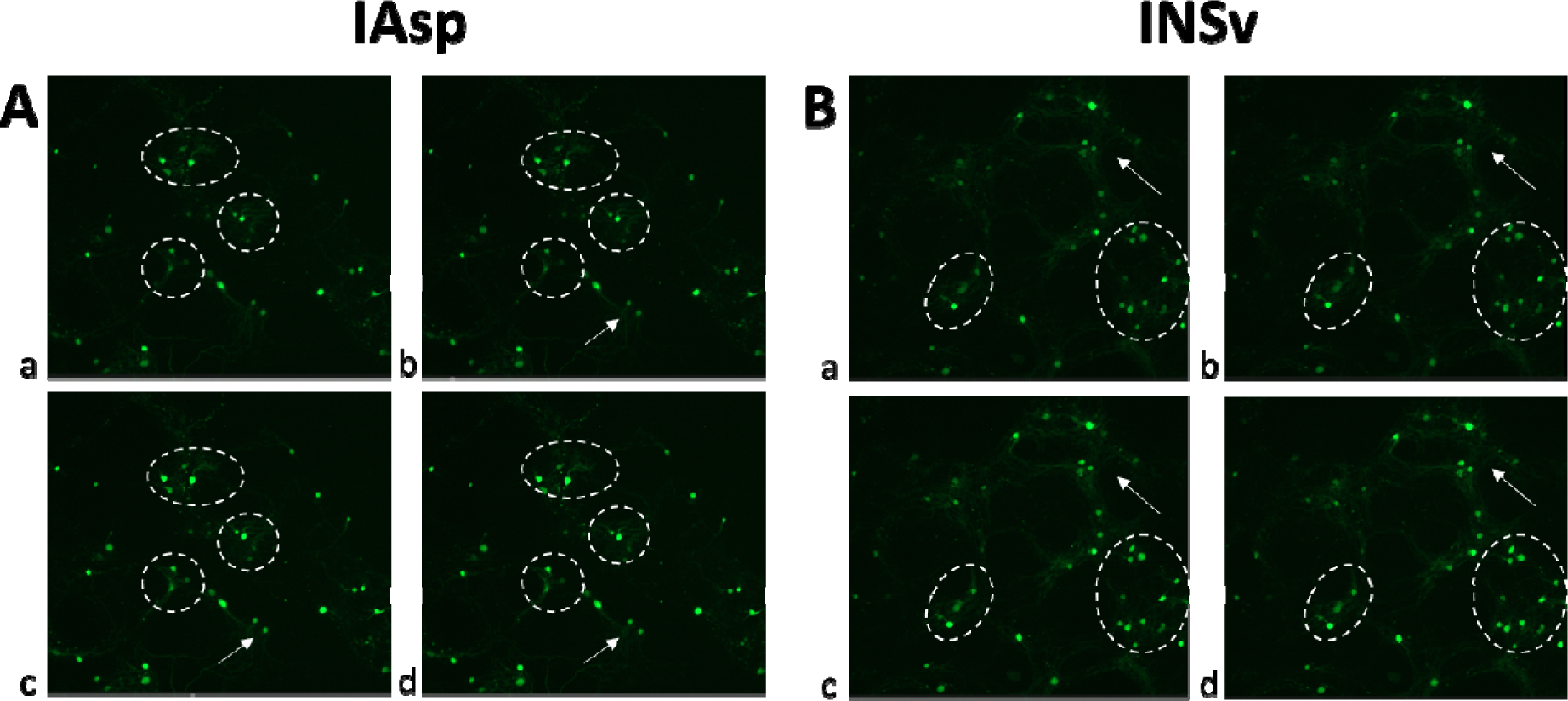
**Selected segments from various recordings of calcium activity associated with IAsp (A) and INSv (B)**. For both panels, a) a pre-stimulation network image and b-d) a progressive spread of calcium activity across the network, with detailed annotations for each stage.

**Supplementary Figure 4.**
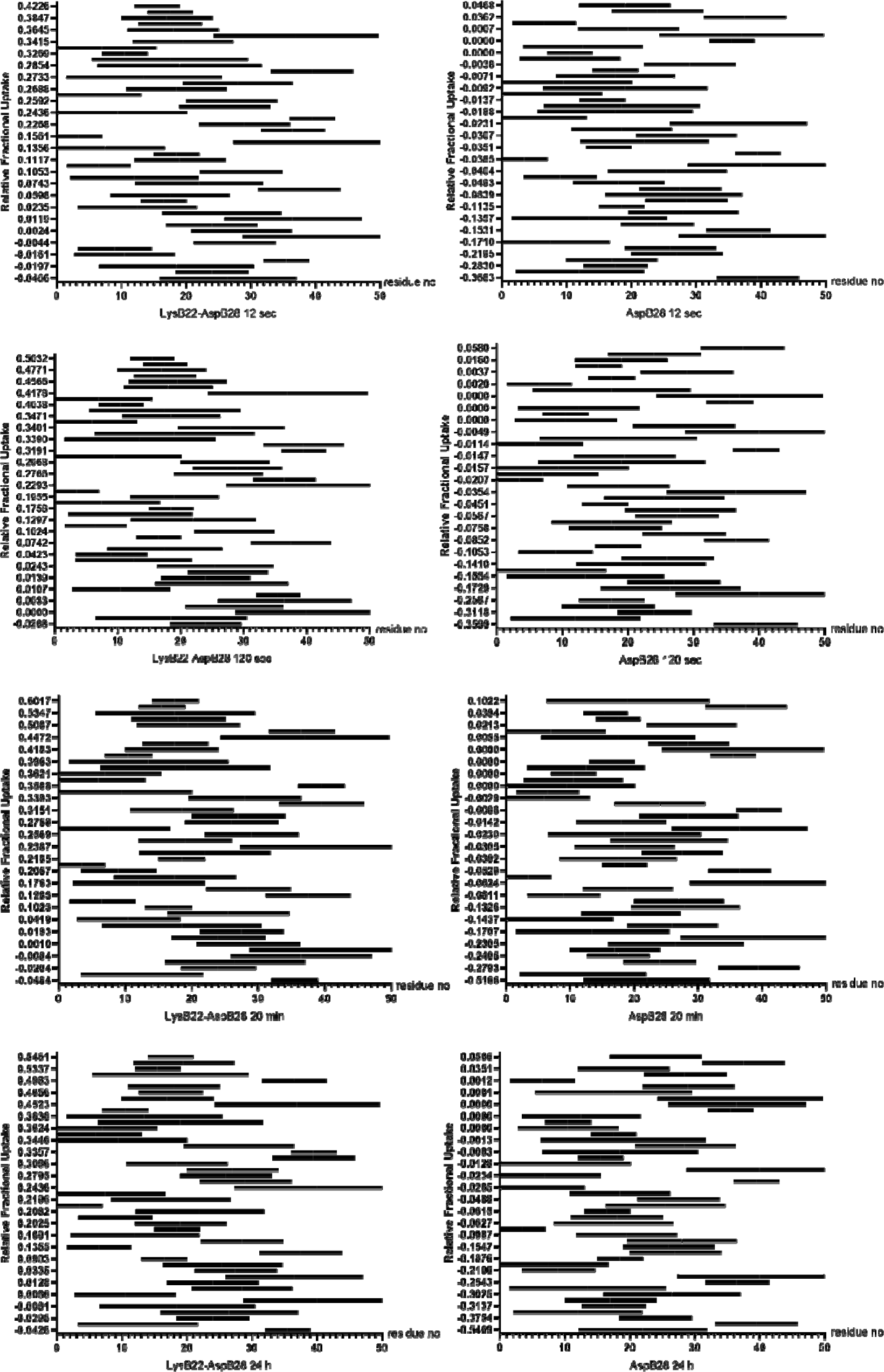
Projection of the peptide-specific deuterium exchange. Relative deuterium exchange of INSv and IAsp peptides depicted by Woods plots over the 12-sec, 120-sec, 20-min, and 24-hour time courses.

**Supplementary Figure 5.**
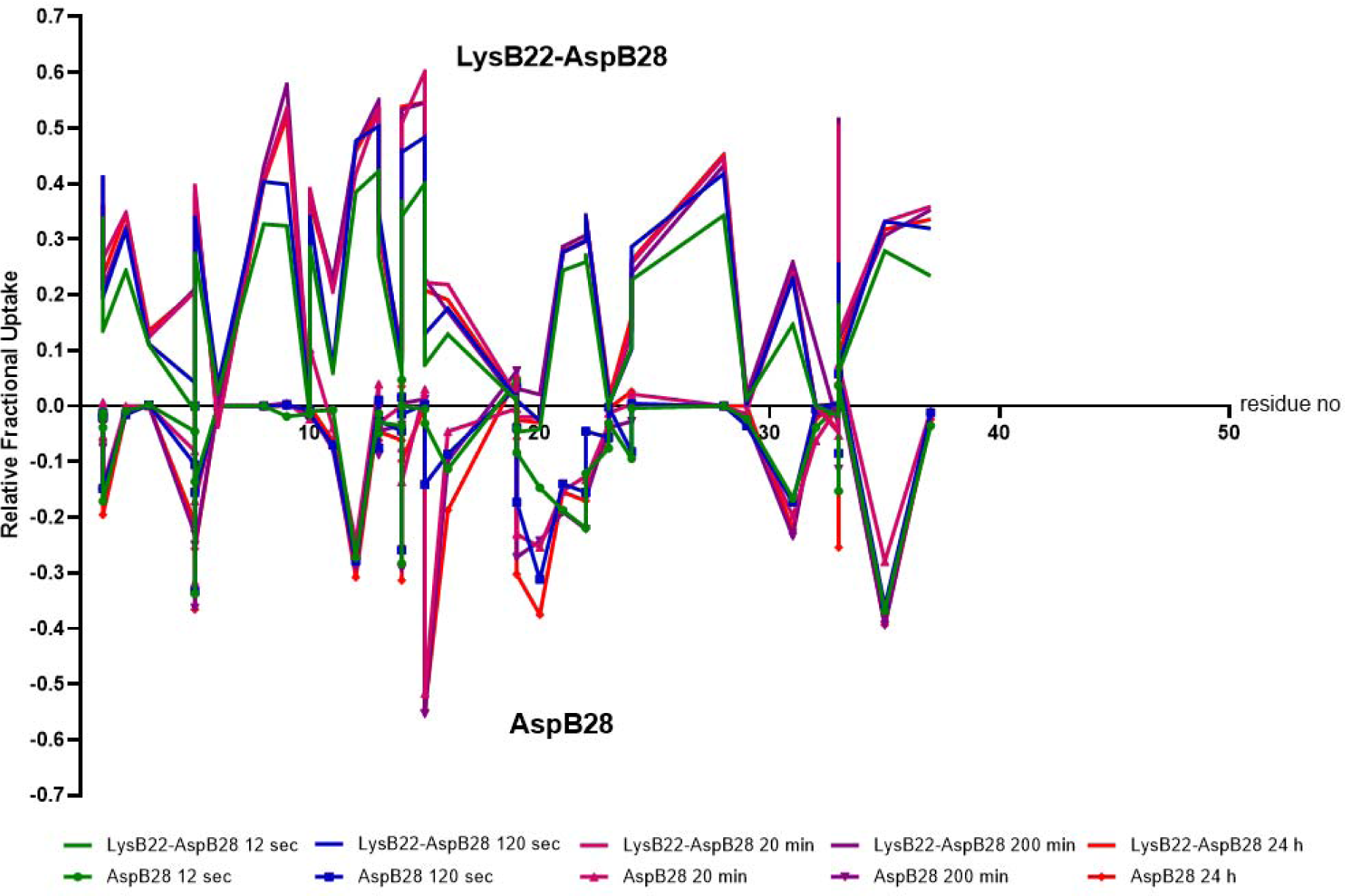
Butterfly plot for deuterium incorporation of INSv (LysB22-AspB28) and IAsp (AspB28). Apart from overall dynamic variability, especially between 20-25 residues in both conformers indicates substantial mobility over all time intervals.

**Supplementary Figure 6.**
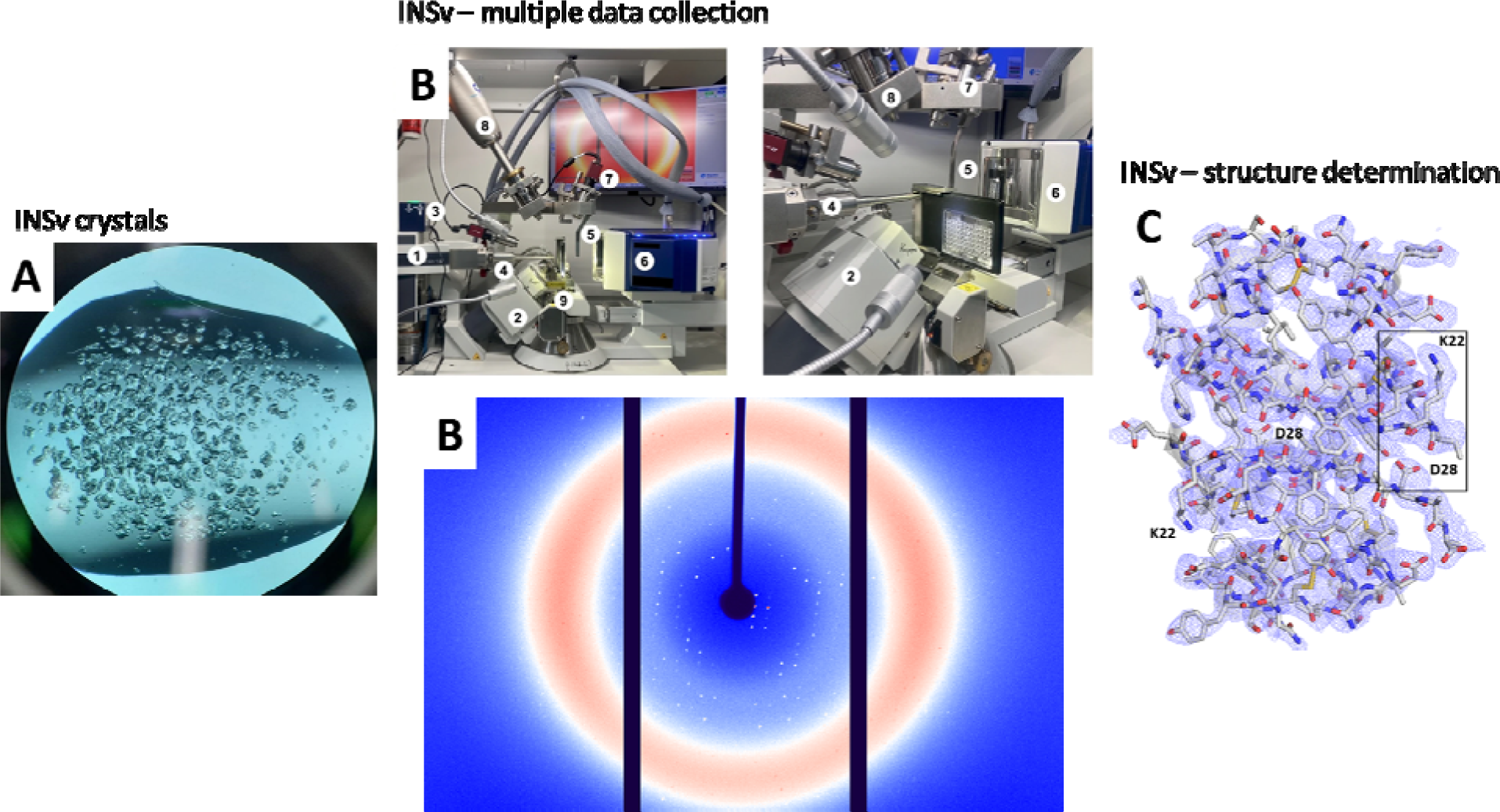
Hexamer INSv data collection and structure determination process. **(A)** INSv hexamers (20 um) has been crystallized in 2.4 M NaCl, 100 mM Tris-HCl at pH 7.4, 6 mM ZnCI2, 20 (w/v) poly(ethylene glycol) PEG-8000 buffer. **(B)** Multiple data collection has been performed by Turkish Light Source at ambient temperature. Main hardware components of Turkish Light Source. (1) X-ray source, (2) four-circle Kappa goniometer, (3) shutter, (4) collimator, (5) beamstop, (6) X-ray detector, (7) video microscope, (8) low-temperature insert, (9) XtalCheck module. (*adapted from Atalay et al.*, *2023; Gul et al.,* 2023). **(C)** Double mutated INSv hexamer has been determined as a dimer in the asymmetric unit at 2.5 A resolution. 2*Fo-Fc* simulated annealing-omit map at 1 sigma level is colored in slate.

